# Arrest-associated transcriptional programmes outweigh apoptosis-associated programmes in treatment-naive pancreatic ductal adenocarcinoma

**DOI:** 10.64898/2026.11.04.764269

**Authors:** Chukwuemeka A. Obuekwe, Blessing T. Ogunoye, Aderinsola Adelaja, Jude Ossai, Adegbesan A. Christopher, Adewunmi Akingbola

**Author notes:** Corresponding author: Chukwuemeka A. Obuekwe, Medical Sciences Division, University of Oxford, Oxford, United Kingdom. Author email addresses: Chukwuemeka A. Obuekwe, Blessing T. Ogunoye, Aderinsola Adelaja, Jude Ossai, Adegbesan A. Christopher, Adewunmi Akingbola.

## Abstract

**Background:** Pancreatic ductal adenocarcinoma (PDAC) is driven by mutant KRAS, and RAS inhibitors now extend survival in previously treated metastatic disease. Whether treatment shrinks a tumour or only halts its growth depends partly on whether cancer cells die or stop dividing, yet how untreated PDAC cells are balanced between these programmes is unknown.

**Aims:** We aimed to characterise the balance between apoptosis-associated and arrest-associated transcriptional programmes in treatment-naive PDAC and to examine their relationships with tumour subtype, TGM2 expression, tissue organisation, survival and drug sensitivity.

**Methods and results:** We scored 52,999 malignant cells from 18 untreated patients in the Hwang atlas using a transcriptional contrast between genes associated with apoptosis and genes associated with resistance to apoptosis or cell-cycle arrest. In every patient the contrast was shifted toward the arrest-associated side: using equality between the two sides as the dividing threshold, 86.3% of cells with an informative score fell on that side, although the proportion depends on the threshold chosen. The shift recurred in the independent Steele atlas (median 71.6% per sample, range 37.1% to 90.8%). In the discovery cohort, tumours with higher basal-like scores had fewer arrest-side cells, and TGM2 expression was modestly higher on the arrest-associated side in both cohorts. The contrast did not track cell-line drug sensitivity or survival in bulk tumours.

**Conclusion:** Untreated PDAC cells show a reproducible transcriptional bias toward anti-apoptotic and cell-cycle-arrest programmes. Whether this bias predicts cellular responses to treatment remains to be tested.

## Introduction

Pancreatic ductal adenocarcinoma (PDAC) remains among the most lethal common cancers, with five-year survival below 15% (Siegel et al., 2024). More than 90% of tumours carry an activating KRAS mutation (Raphael et al., 2017), and drugs that block RAS signalling are now reaching patients: in the phase 3 RASolute 302 trial, the RAS(ON) inhibitor daraxonrasib extended overall survival compared with chemotherapy in previously treated metastatic PDAC (13.2 versus 6.7 months; O’Reilly et al., 2026). Whether such a drug shrinks a tumour or only halts its growth depends in part on what happens to individual cancer cells when RAS signalling is blocked: some die by apoptosis, while others stop dividing and survive.

Which of these outcomes a cell follows is set partly by its state before treatment, including the balance of pro- and anti-apoptotic BCL-2 family proteins and the activity of cell-cycle brakes, although gene expression alone cannot show what a given cell will do. Single-cell and transcriptomic studies have mapped extensive heterogeneity among PDAC cells (Hwang et al., 2022; Chan-Seng-Yue et al., 2020), and preclinical work links cell state to response: a classical epithelial state confers acute resistance to KRAS inhibition (Singhal et al., 2024), whereas basal-like and mesenchymal states in mouse models respond more favourably (Dilly et al., 2024). How untreated human PDAC cells are balanced between apoptosis- and arrest-associated programmes has not, to our knowledge, been measured directly.

Here we measure that balance cell by cell. We combine curated gene modules into a single contrast that compares pro-apoptotic and caspase/mitochondrial genes with anti-apoptotic and cell-cycle-arrest genes, and apply it to treatment-naive tumours from two independent single-cell atlases. We then relate it to tumour subtype, identify a gene that tracks it, and test whether it extends to tissue architecture, survival and drug sensitivity. We use cytostatic-prone only as a shorthand for cells on the arrest-associated side of this contrast; it describes expression, not an observed or predicted response to treatment.

## Materials and Methods

### Software and reproducibility

All analyses were performed in R 4.6.0 on macOS. Principal packages were UCell 2.16.0, Seurat 5.5.1 with SeuratObject 5.4.0, harmony 2.0.5, DESeq2 1.52.0, fgsea 1.38.0, GSVA 2.6.3, LIANA 0.1.14, survival 3.8.9, survminer 0.5.2, ggplot2 4.0.3, patchwork 1.3.2, Matrix 1.7.5 and hdf5r 1.3.12. The R scripts, project environment (including the renv lockfile), workflow and supporting processed objects are available in the public repository cited in the Data Availability Statement. The scripts reproduce the reported statistics from the deposited processed objects; the original source datasets are not mirrored in the repository and should be obtained from the public repositories and accessions listed below. The repository README documents data retrieval, the analysis run order, and the mapping between scripts and manuscript figures and tables. It also identifies a small number of figure panels that were generated separately from an intermediate object too large to deposit. Where a reported value depends on an analytical choice, that choice is stated alongside it in Results.

### Public datasets

GSE202051 provided the primary single-nucleus atlas of human PDAC (Hwang et al., 2022): 44 samples from 43 patients, of which 18 untreated patients contributed one sample each. GSE155698 provided the replication single-cell atlas (Steele et al., 2020). GSE272362 provided Visium spatial transcriptomics of human PDAC (Khaliq et al., 2024), of which the ten primary tumour sections were analysed. TCGA PAAD STAR gene-level quantifications for 178 primary tumour samples were obtained from the NCI Genomic Data Commons with TCGAbiolinks (Colaprico et al., 2016), overall survival from the clinical annotations curated for the TCGA PAAD marker paper (Raphael et al., 2017), and ABSOLUTE tumour purity estimates from the PanCanAtlas consolidated purity and ploidy table (TCGA_mastercalls.abs_tables_JSedit.fixed.txt), which applies the ABSOLUTE method (Carter et al., 2012). DepMap 24Q2 (Broad Institute) provided CRISPR-Cas9 Chronos gene effect scores and baseline log2(TPM + 1) expression across cancer cell lines (Meyers et al., 2017; Dempster et al., 2021). GSE269985 provided bulk transcriptomes of parental and RAS inhibitor-resistant PDAC cell lines (Dilly et al., 2024).

### Gene module construction

Three exploratory axes accompany the primary apoptosis/arrest contrast (Axis 2); their results are presented mainly in the Supplementary Information. Axis 1 comprises two modules. The MAPK output module (DUSP4, DUSP6, ETV4, SPRY4, EGR1, FOSL1, CCND1, SPRED1) is a curated set of canonical ERK-MAPK transcriptional targets, selected a priori from pathway biology and not from any correlation with KRAS dependency. The bypass potential module (EGFR, AXL, FGFR1, FGFR3, ETV1, ETV5, PIK3CA, MTOR) comprises receptor tyrosine kinase and downstream signalling genes selected as candidate bypass effectors and retained because each was significantly upregulated in the Panc 02.03 resistance contrast described below (FDR-adjusted p < 0.05; log2 fold changes 0.20 to 2.87). Only two of the eight, FGFR1 and FGFR3, were also significantly upregulated in the PANC-1 contrast, so this module rests predominantly on evidence from a single cell line.

A separate benchmarking module, DepMap correlates, not part of the Axis 1 calculation, comprises the 30 genes whose expression correlated most negatively with KRAS Chronos dependency across the 44-line PDAC panel (Pearson r from -0.565 to -0.466, from 131 genes with r < -0.4 among 18,784 tested). Because these genes were selected using the same metric against which the module is subsequently assessed, this module is circular by construction. It is reported for comparison only and is not used as validating evidence.

Axis 2 comprises four curated modules: anti-apoptotic (BCL2, BCL2A1, BCL2L1, BCL2L2, MCL1), pro-apoptotic (BAD, BID, BIK, BMF, BBC3, PMAIP1, BCL2L11), caspase and mitochondrial apoptosis-associated (CASP3, CASP7, CASP8, CASP9, CYCS, DIABLO) and cytostasis (CDKN1A, CDKN2A, RB1, GADD45A, GADD45B). These gene sets were specified from BCL-2 family, caspase cascade and cell-cycle-arrest biology rather than selected by differential expression; the behaviour of their constituent genes in the resistance contrast is reported in Results as evidence bearing on the axis, not as the basis for membership.

Axis 3 and Axis 4 modules are the leading-edge gene sets of Hallmark pathways significantly enriched in the resistance contrast (fgsea, 1,000 permutations, minimum set size 15). Axis 3 comprises the hypoxia leading edge (48 genes; normalised enrichment score 1.67) and the oxidative phosphorylation leading edge (87 genes; NES -1.66). Axis 4 comprises the union of leading-edge genes from five immune-related sets (interferon alpha, interferon gamma, TNF-alpha signalling via NF-kB, IL6-JAK-STAT3 and inflammatory response; 157 genes), with a core module of the 66 genes appearing in two or more of them.

### Resistance transcriptomics

The resistance transcriptomic analysis was restricted to the two human PDAC lines in GSE269985, Panc 02.03 and PANC-1, so that it remained in human cells like the rest of the study; the murine KPC-derived lines in the same series were not included. Each human line contributed three parental and three resistant replicates (n = 12). Differential expression between resistant and parental cells used DESeq2 1.52.0 with design ∼cell_line + condition + cell_line:condition (Love et al., 2014). The condition coefficient gives the resistance effect at the Panc 02.03 reference level; the PANC-1 effect was obtained by combining the main effect with the interaction term. A gene was considered cross-line concordant if it reached FDR-adjusted significance in both line-specific contrasts with the same direction of effect. Supplementary Fig. 3 displays the Panc 02.03 contrast with shrunken fold-change estimates; it is a single-line result and is labelled as such.

### Single-cell scoring and classification

GSE202051 was read from its h5ad container using hdf5r (Hoefling and Annau, 2021). Malignant cells were taken from the published annotation, giving a 64,538 by 22,164 matrix of log-normalised values. All 11 modules were scored with UCell (maxRank 1500) (Andreatta and Carmona, 2021, 2026).

Axis 1 was computed per cell as log2((MAPK output + ε) / (bypass potential + ε)) and Axis 2 as log2((pro-apoptotic + caspase/mitochondrial + ε) / (anti-apoptotic + cytostasis + ε)), with ε = 0.001. For Axis 2, the two modules on each side are summed so that each side of the contrast is represented by its full gene complement, and the log ratio makes the score symmetric about zero: it is zero when the two composites are equal, negative when anti-apoptotic and cytostasis signal dominates, and approximately plus or minus one when either side is twice the other. The constant ε is a pseudocount, a small value added to both numerator and denominator so that the ratio remains defined when a composite is zero, which UCell returns when no gene of a module ranks within the cutoff. Its value cannot change which side of zero a cell falls on: for any positive ε, the ratio exceeds one exactly when the apoptosis-associated composite exceeds the arrest-associated composite. Every classification at the zero reference, including all prevalence estimates, is therefore independent of the value chosen, and a simple difference between the composites would classify cells identically, so the two forms are not independent checks of each other. The value of ε affects only the magnitude of the score, chiefly in cells where one composite is close to zero, and so bears on the shape of the continuous distribution and on thresholds other than zero. Reconstruction of both formulas from the stored per-cell module scores reproduced the saved values exactly. Axes 3 and 4 were used as continuous UCell scores.

Cells with an Axis 2 score of exactly zero were classified as uninformative and excluded from prevalence estimates, an exclusion defined after the initial analysis rather than specified in advance. A score of exactly zero means that the two composites are equal; checking the stored module scores showed that in every such cell all four module scores were zero, with no cell scoring zero through equal non-zero sums, and the smallest absolute score among the remaining cells was 0.00057. Their number and causes are reported in Results. Among informative cells, those with Axis 2 below zero were classified as being on the arrest-associated side of the contrast (cytostatic-prone) and those above zero on the apoptosis-associated side; these are labels for positions on a transcriptional contrast, not measured cellular fates. A median split of Axis 1 provided an orthogonal sub-classification into MAPK-output-high and bypass-signature-high cells, giving four quadrants in combination with the Axis 2 split. Axis 2 alone defines which side of the contrast a cell falls on. Sensitivity to the threshold was assessed by repeating the classification with a median split and a bottom-tertile split of the Axis 2 distribution, and by sweeping the threshold across its range. The sweep is reported both as the pooled proportion of informative cells and as the number of patients in which cytostatic-prone cells exceed half, because these two criteria diverge. Sensitivity to the UCell ranking cutoff was assessed separately by recalculating all four Axis 2 module scores at maxRank values of 1,000, 1,500, 2,000 and 2,500 and repeating the classification from the recalculated scores; results are reported with both the informative-cell and the whole-cohort denominator (Supplementary Table 1).

Specificity was assessed by permutation against random gene sets drawn from expressed genes and matched to the real modules by detection-frequency decile, with the four module sizes preserved, on a random subsample of 8,000 untreated cells. In this analysis, random gene sets preserving the unequal numerator and denominator sizes produced a negatively skewed null distribution. Specificity was therefore evaluated against an additional null with equalised module sizes, obtained by randomly subsampling the two larger modules to five genes each, applying the same scoring and classification procedure to the observed and random gene sets. One thousand permutations were used for the size-equalised null. The test statistic is the percentage of informative cells classified cytostatic-prone. Because equalisation requires randomly subsampling the two larger modules to five genes each, the observed statistic is not a single value: it was recomputed across 100 independent equalisation draws, and the reported observed value is the median of that distribution, whose full range is also reported. Each permutation applies the identical procedure to detection-matched random genes, drawing all four modules at five genes each, so the equalisation step is handled the same way in the observed and null arms. The empirical p-value is the proportion of permutations reaching or exceeding the observed median, with an add-one correction, and null intervals are reported as 2.5th to 97.5th percentiles. Sensitivity to the cell subsample was checked by repeating the classification on three independent 8,000-cell subsamples and on all informative cells, and by repeating the whole permutation procedure on two further independent subsamples (500 permutations each). Seeds are set in the deposited scripts.

Two analytical decisions were specified before scoring: restriction of primary analysis to treatment-naive cells with treated cells analysed separately, and a merge threshold of Pearson r > 0.6 for combining Axes 3 and 4, which was not met.

For visualisation only, log-normalised counts were processed in Seurat with Harmony batch correction across patients, and UMAP computed on Harmony dimensions 1 to 20. Malignant cells remain partly separated by patient after integration, as expected for clonal tumour transcriptomes (Supplementary Fig. 9a). No reported statistic depends on the embedding, and all prevalence results are summarised per patient.

Moffitt classical and basal-like signatures (Moffitt et al., 2015) were scored per cell with UCell. Patients were assigned a dominant subtype by comparing mean scores. Patient-level association between the cytostatic-prone proportion among informative cells and the mean basal-like score, averaged across all untreated malignant cells of a patient, used an exact Spearman test across the 18 untreated patients. Uncertainty was estimated by percentile bootstrap over patients (10,000 resamples) and influence by leave-one-out. Because per-patient prevalence could reflect data quality, the association was additionally adjusted for the per-patient fraction of uninformative cells and for mean genes detected, using partial Spearman correlation implemented as the correlation of rank residuals. The replication atlas was scored with identical modules, formulas and thresholds, including identical handling of uninformative cells. For the TGM2 comparison in that atlas, per-sample filtered count matrices were obtained from the individual sample records and TGM2 counts normalised per cell to counts per 10,000 followed by a log transformation, matching the normalisation of the discovery matrix. Cells were matched to the existing classifications by sample and barcode, and all scored cells were recovered. The classification was not recomputed, and TGM2 contributes to none of the gene modules, so the comparison is independent of the classification.

### Ligand-receptor inference

LIANA 0.1.14 (Dimitrov et al., 2022) was applied to untreated cells subsampled to at most 500 per annotated cell type (16,090 cells), using OmniPath as the prior resource (Turei et al., 2021) and four scoring methods (NATMI, SingleCellSignalR, Connectome, logFC) combined by rank aggregation. Interactions were compared between malignant senders on the arrest-associated and apoptosis-associated sides of the contrast using the aggregate rank score, which is a rank-derived probability and not a measure of interaction strength; ratios of these scores are reported as differences in ranking only. TGM2 and ITGA4 expression were compared between states using the stored log-normalised values, with per-patient means tested by a paired Wilcoxon signed-rank test across the 18 untreated patients, so that cells are not treated as independent. Confidence intervals for the mean paired difference are conventional t-based intervals on the per-patient differences, in both the discovery and the replication cohort; the standardised paired effect is the mean of the per-patient differences divided by their standard deviation.

### Spatial analysis

The GSE272362 Seurat object supplies both RCTD cell type assignments (dominant type and spot class) and a precomputed fges assay of functional signature scores; both were used as supplied and were not recomputed here. Analysis used the Treg activation, checkpoint molecule, myeloid suppression and effector cell scores, the last as a specificity control. For each primary PDAC section, Euclidean distance from every non-epithelial spot to the nearest tumour epithelial spot was computed, and the Spearman correlation between distance and each programme score obtained per section. Because Visium spots are spatially autocorrelated, spot-level p-values were not used for inference; instead inference was based on the per-section correlation coefficients. Sections with fewer than ten epithelial anchor spots were excluded, leaving nine; the excluded section contained nine anchor spots, against at least 99 in every other section, too few to define a distance gradient, and its inclusion is reported as a sensitivity check. These coefficients were tested with an exact two-sided Wilcoxon signed-rank test, and Benjamini-Hochberg adjustment was applied across the four programmes within each analysis. Specificity was assessed by correlating each section’s effector coefficient with its Treg coefficient across sections. A sensitivity analysis repeated the correlations in the same nine sections using only non-epithelial spots that RCTD classified as confident singlets, with epithelial anchors defined as in the primary analysis; per-section coefficients were compared with the all-spot estimates by paired Wilcoxon signed-rank test.

### Bulk tumour analyses

Expression for the 178 primary tumour samples was transformed to log2(TPM + 1), and every module of the four axes was scored with ssGSEA (GSVA, default normalisation). The bulk Axis 2 contrast was calculated from ssGSEA scores: half the difference between the summed apoptosis-associated module scores (pro-apoptotic and caspase/mitochondrial) and the summed arrest-associated module scores (anti-apoptotic and cytostasis), so that lower values indicate relatively greater arrest-associated signal. Rescoring the raw files reproduced the stored bulk scores (Pearson r > 0.9999) and assigned every sample to the same median-split group. Survival analyses used the 146 samples with complete overall survival data, and Kaplan-Meier curves and Cox proportional hazards models used a median split of the bulk Axis 2 score (73 per group). High cytostatic, meaning the lower Axis 2 score, is the reference level, so the reported hazard ratio is that of the low cytostatic group relative to it; a hazard ratio above one therefore corresponds to shorter survival in the low cytostatic group, consistent with the Kaplan-Meier medians. Both the Cox Wald p-value and the log-rank p-value are reported, each labelled. Association between the Axis 2 score and ABSOLUTE purity was assessed by Spearman correlation in the 133 samples carrying a purity estimate, matched on the first 15 characters of the TCGA barcode. All ABSOLUTE metrics in that table (purity, ploidy, cancer DNA fraction, subclonal genome fraction and genome doublings) were tested against all four axes.

### Cell-line analyses

For cell lines the apoptosis/arrest contrast was computed from bulk expression rather than with UCell, whose rank-based scoring is designed for sparse single-cell data. Each of the four modules was summarised as the mean log2(TPM + 1) expression of its member genes present in the DepMap expression matrix, and the cell-line apoptosis/arrest contrast was defined as the mean of the pro-apoptotic and caspase/mitochondrial module means minus the mean of the anti-apoptotic and cytostasis module means. Positive values indicate relatively greater apoptosis-associated expression and negative values relatively greater arrest-associated expression. This cell-line contrast is related to, but distinct from, the single-cell UCell log ratio, and it underlies all cell-line analyses reported here. Absolute contrast scores are therefore not directly comparable between single cells, bulk tumours and cell lines; across the 44 lines it correlated with an ssGSEA-based alternative at Spearman r = 0.96. The 44 lines were the PDAC lines with both expression and CRISPR data, defined as the intersection of 51 lines with expression (after removing duplicate model identifiers) and 46 with essentiality data. Lines were split at the median contrast of this panel: the 22 lines with lower scores were classified as high-cytostatic and the 22 with higher scores as low-cytostatic. Chronos scores for TGM2, NT5E and KRAS were compared between groups by Wilcoxon rank-sum test.

Drug sensitivity was taken from GDSC2 release 8.5 (fitted dose response, October 2023). The 44 lines were matched to GDSC cell lines by stripped cell line name from the DepMap model metadata, with COSMIC identifier as a fallback; 24 matched. Where a line had been screened more than once for a compound, values were averaged. Sensitivity was related to the cell-line contrast by Spearman correlation, with the prediction that higher scores, indicating relatively greater apoptosis-associated expression, would accompany lower ln(IC50), giving a negative correlation. Two tests were run in sequence and both are reported. In the first, six MAPK-pathway inhibitors were designated the primary drug set before any association was computed, with Benjamini-Hochberg adjustment across the set; compounds outside it are reported only as context, alongside the number of nominally significant results expected by chance. After the first analysis, a second test was specified because the contrast includes BCL-2 family genes. Its expected direction and drug sets were fixed before the second set of associations was calculated: nine BH3 mimetics and direct apoptosis modulators as the primary set, with Benjamini-Hochberg adjustment within it, and nine cytotoxic agents as a secondary set reported without adjustment, with the earlier MAPK set retained as a reference. ln(IC50) was the primary metric and area under the dose-response curve a secondary one.

## Results

### A transcriptional contrast between apoptosis- and arrest-associated programmes

To measure this balance we built a contrast from four curated gene modules. On the apoptosis-associated side are pro-apoptotic BH3-only genes (BAD, BID, BIK, BMF, BBC3, PMAIP1, BCL2L11) and caspase and mitochondrial genes (CASP3, CASP7, CASP8, CASP9, CYCS, DIABLO); on the arrest-associated side are anti-apoptotic BCL-2 family genes (BCL2, BCL2A1, BCL2L1, BCL2L2, MCL1) and cell-cycle-arrest genes (CDKN1A, CDKN2A, RB1, GADD45A, GADD45B). The genes were chosen from established apoptosis and cell-cycle biology, not selected from the data. Each module is scored in every cell, and the two sides are compared as a log ratio. Negative values indicate relatively greater anti-apoptotic and arrest-associated expression; positive values indicate relatively greater apoptosis-associated expression (Methods). We call this contrast Axis 2 because it was developed alongside three exploratory axes for RAS signalling balance, hypoxia-metabolic state and innate immune mimicry, which are described in the Supplementary Information (Supplementary Figs 1 and 2).

Expression changes in PDAC cell lines with acquired resistance to RAS inhibition prompted the choice of these modules but did not define them, and the changes were not consistent between the two resistant lines: only BCL2, PMAIP1, RB1 and GADD45A (up) and BIRC5 and GADD45B (down) changed significantly in the same direction in both, while CDKN1A moved in opposite directions (+1.04 in Panc 02.03, -0.61 in PANC-1; Supplementary Figs 3 and 4).

### The contrast is shifted toward arrest-associated programmes in every patient

We scored 64,538 malignant cells from the Hwang et al. single-nucleus atlas: 52,999 from 18 treatment-naive patients and 11,539 from 25 patients who had received neoadjuvant therapy (Hwang et al., 2022). In 7,825 treatment-naive cells (14.8%) no module registered at all, either because no module gene was detected (6,217 cells) or because the few detected genes were tied at low expression with many others and so ranked outside the scoring cutoff (1,608 cells). These cells had insufficient detectable expression of the selected genes to determine which programme predominated. They also had fewer genes detected overall (median 993 versus 1,549) and were excluded from prevalence estimates (Methods; shown in grey in Supplementary Fig. 5).

In all 18 patients the distribution of scores was shifted toward the arrest-associated side (Fig. 1a). Taking the point where the two sides are equal, a score of zero, as the dividing line, 86.3% of the 45,174 scored cells were on the arrest-associated side, and the per-patient figure ranged from 69.0% to 95.1% (median 87.7%; Fig. 1b). This dividing line follows from the definition of the score rather than from the data, and the percentage depends on where it is placed: dividing instead at the median or at the lowest third of the pooled scores gives per-patient proportions of 28.5% to 61.9% and 14.9% to 52.2% (Supplementary Fig. 6). Across all scored cells the arrest-associated side remains in the majority for any dividing line above -6.0, and within every patient for any line above -1.0. The robust finding is therefore the direction and consistency of the shift, not a particular percentage. The 86.3% describes the proportion below zero on this transcriptional score; it does not mean that 86.3% of cells would arrest rather than undergo apoptosis after treatment.

**Figure 1.**
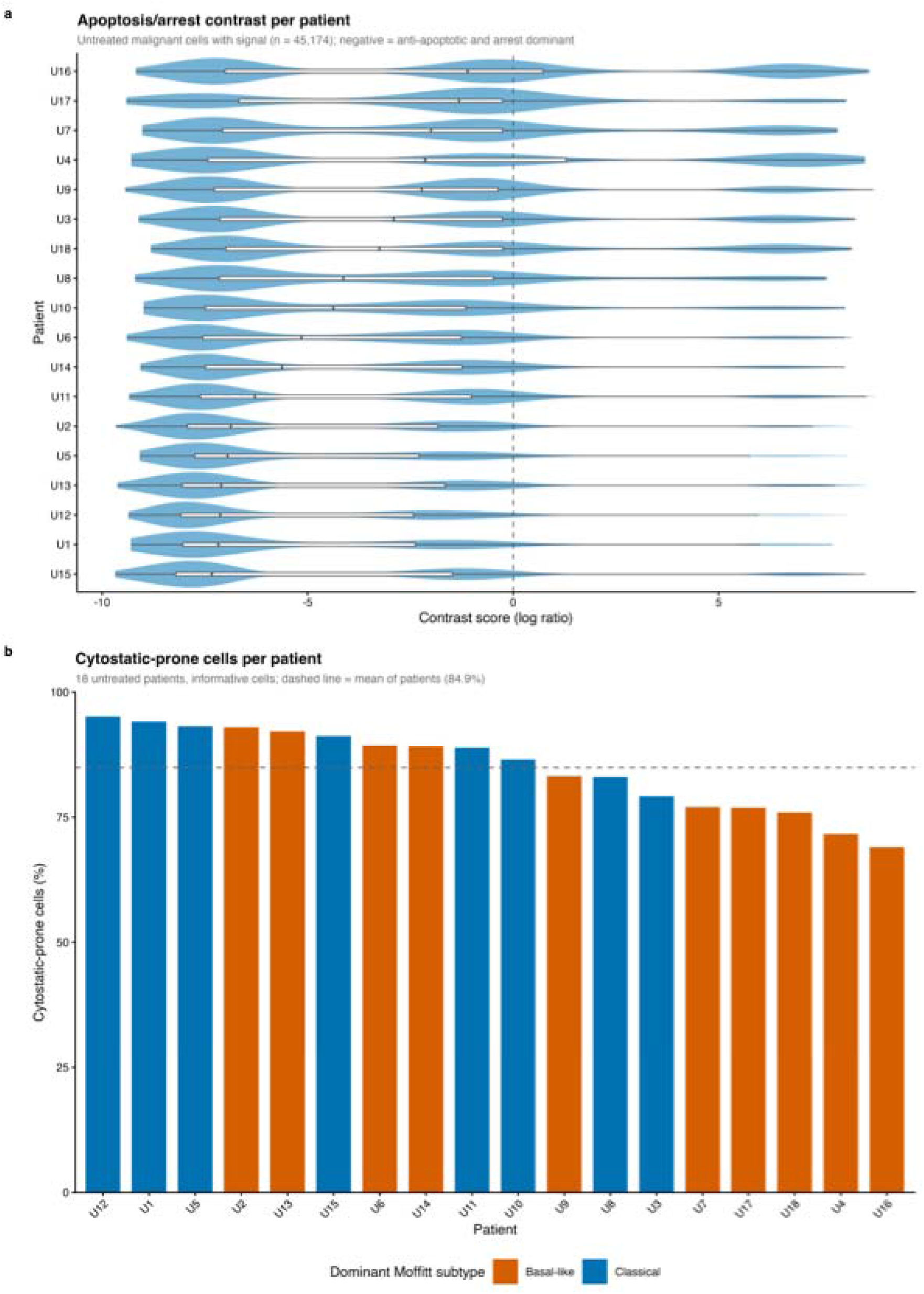
Distribution of the apoptosis/arrest transcriptional contrast in treatment-naive PDAC. a) Per-cell contrast score for each of the 18 treatment-naive patients, ordered by median; negative values indicate dominance of anti-apoptotic and arrest-associated signal. Dashed line marks a score of zero, where the two sides are equal. Cells with no measurable score are excluded. b) Proportion of scored cells on the arrest-associated side per patient, coloured by the patient’s dominant Moffitt subtype; dashed line marks the mean of patient proportions.

The shift was not an artefact of the scoring procedure or of the genes chosen. Changing the scoring cutoff from 1,000 to 2,500 genes altered the overall proportion only between 86.0% and 86.9%, with every patient above half at each setting, although the share of cells with a measurable score varied from 76.0% to 88.1% (Supplementary Table 1). Random gene sets matched to the real modules for size and detection placed a median of 49.2% of cells on the arrest-associated side, against 87.9% for the real modules; four of 1,000 random sets reached the observed value (p = 0.005), and the result held in two further independent subsamples (Supplementary Table 2). The shift is therefore not a property of arbitrary genes, although this alone does not establish biological specificity. Tumours from patients who had received neoadjuvant therapy showed a similar shift (82.1%), but these are different patients sampled once, so no effect of treatment can be inferred.

### The shift recurs in an independent atlas and varies with tumour subtype

The shift recurred in an independent atlas generated on a different platform (Steele et al., 2020). Scored in exactly the same way, 13.3% of 14,571 malignant cells had no measurable score, and the median proportion on the arrest-associated side was 71.6% per sample (range 37.1% to 90.8%; Fig. 2b). The direction was reproduced at the level of the cohort but not in every sample: all 18 Hwang patients exceeded half, whereas some Steele samples did not.

**Figure 2.**
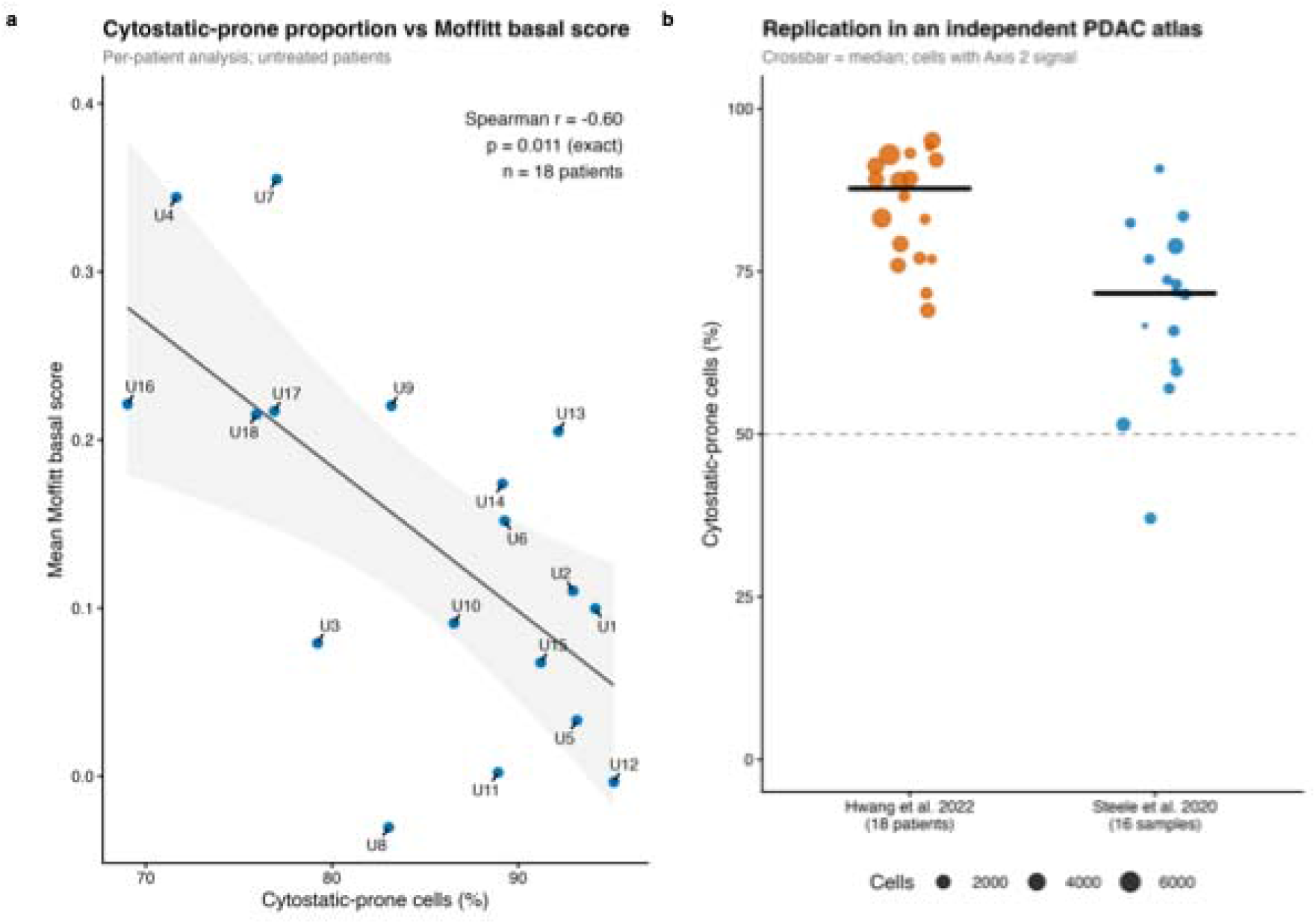
Subtype association and independent replication. a) Patient-level relationship between the proportion of informative cells on the arrest-associated side and the mean Moffitt basal-like score across all untreated malignant cells of each patient (exact Spearman test, n = 18). b) Per-sample proportions in the Hwang and Steele atlases, scored with identical modules, formulas and thresholds including identical exclusion of uninformative cells; crossbars mark medians and point size indicates cells per sample.

The shift was weaker in tumours of more basal-like subtype. Tumours with higher mean Moffitt basal-like scores contained fewer arrest-side cells (Spearman r = -0.60, 95% bootstrap confidence interval -0.85 to -0.13, p = 0.011, n = 18; leave-one-out range -0.73 to - 0.55; Fig. 2a). Differences in data quality between tumours did not explain this, because the association persisted after adjustment for each patient’s fraction of unscored cells (partial r = -0.52, p = 0.027) or mean genes detected (partial r = -0.54, p = 0.022). Within tumours, by contrast, cells with a higher basal-like than classical score and those with a higher classical than basal-like score were on the arrest-associated side at almost the same rate (86.1% versus 86.6%), so the relationship operates between tumours rather than between individual cells. Its direction agrees with preclinical reports that basal-like and mesenchymal states respond better to KRAS-pathway inhibition (Dilly et al., 2024; Singhal et al., 2024), but it rests on 18 patients and a wide confidence interval.

### TGM2 is a reproducible correlate of the arrest-associated side

To explore genes associated with the arrest-associated side, we compared computationally inferred ligand-receptor interactions between malignant cells on each side in a 16,090-cell subsample (Dimitrov et al., 2022). A predicted interaction between TGM2 (encoding transglutaminase 2, or TG2) on tumour cells and ITGA4 on immune cells ranked higher when arrest-side cells were the senders, across every ITGA4-expressing immune population tested (Fig. 3b). These rankings indicate relative priority, not interaction strength.

**Figure 3.**
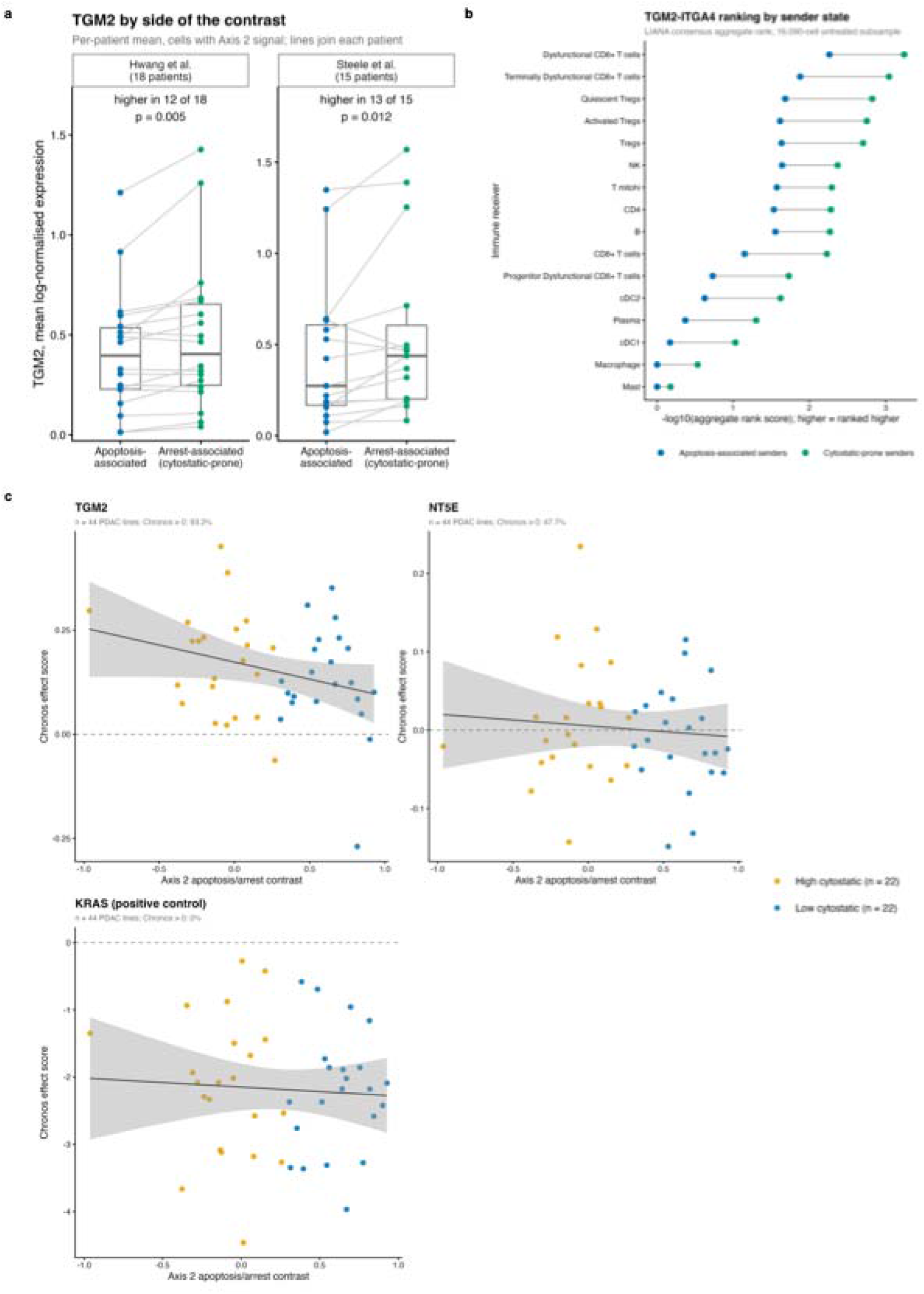
TGM2 as a reproducible correlate. a) Mean TGM2 expression per sample in cells on each side of the contrast, with lines joining each sample, for the discovery cohort (Hwang et al.; 18 patients) and the replication cohort (Steele et al.; 15 patients, two specimens from one patient pooled). Paired Wilcoxon signed-rank test within each cohort. b) LIANA consensus aggregate rank scores for the predicted TGM2-ITGA4 interaction with each immune receiver, shown separately for senders on each side of the contrast; higher values indicate a higher-ranked interaction. These are rank-derived scores, not measures of interaction strength. c) DepMap Chronos gene effect against the cell-line apoptosis/arrest contrast across 44 PDAC cell lines for TGM2, NT5E and KRAS as a positive control; dashed line marks Chronos = 0.

Because TGM2 was nominated by this screen rather than tested in advance, we measured its expression directly in both cohorts. In the discovery cohort TGM2 was detected in 28.6% of arrest-side cells and 21.0% of apoptosis-side cells, and its mean expression was higher on the arrest-associated side in 12 of 18 patients (mean paired difference 0.076 log-normalised units, t-based 95% CI 0.024 to 0.128; paired Wilcoxon p = 0.005). In the replication cohort, where all 14,571 malignant cells included in scoring were matched to their TGM2 counts, the same held in 13 of 15 patients (with two specimens from one patient pooled) (mean paired difference 0.114 log-normalised units, t-based 95% CI 0.013 to 0.216; paired Wilcoxon p = 0.012; Fig. 3a). TGM2 is not one of the module genes, so this comparison is independent of the score.

TGM2 knockout did not impair growth in 93.2% of the 44 DepMap PDAC cell lines (Chronos score above zero; Fig. 3c), so it is not a cell-intrinsic dependency in culture. The association is reproducible but modest, and these data do not show that TGM2 contributes to the arrest-associated state or to any immune effect.

### Tissue organisation, prognosis and drug response

We next examined related questions beyond the single-cell data: immune organisation in tissue, survival in bulk tumours and drug sensitivity in cell lines. The spatial analysis was motivated by reported tumour–immune functions of TGM2; it did not directly map cells classified by the apoptosis/arrest contrast.

Immune programmes in tissue were not concentrated around tumour cells. In nine spatial transcriptomic sections of primary PDAC (Khaliq et al., 2024), scores for Treg activation, checkpoint molecules and myeloid suppression rose, rather than fell, with distance from tumour epithelium, and an effector T cell programme included as a control rose in the same way (median per-section Spearman r 0.153, 0.150, 0.119 and 0.185; positive in 8, 8, 9 and 8 of 9 sections; Wilcoxon p = 0.0078, 0.0078, 0.0039 and 0.0078, each 0.0078 after Benjamini-Hochberg adjustment across the four programmes; Supplementary Fig. 8a, 8b). The effector and Treg gradients were almost perfectly correlated across sections (Spearman r = 0.983). The pattern was unchanged when only spots assigned to a single cell type were used (adjusted p = 0.0052 to 0.0078; effector against Treg r = 0.933; no detectable difference from all spots, paired Wilcoxon p ≥ 0.57) and when the one section with too few tumour spots was included (adjusted p = 0.0098 for all four programmes). The parallel gradients of suppressive and effector programmes are more consistent with general immune compartmentalisation than with selective enrichment of suppressive programmes near tumour epithelium (Supplementary Fig. 7).

The contrast did not predict survival in bulk tumours. In TCGA PAAD (n = 146), patients below the median score had longer median overall survival (691 versus 511 days), but the difference was not statistically significant (hazard ratio 1.386 for the higher-score group relative to the lower-score group, 95% confidence interval 0.874 to 2.197; Cox Wald p = 0.165, log-rank p = 0.164; Supplementary Fig. 8c). The score was only weakly related to tumour purity (Spearman r = 0.153, p = 0.078, n = 133; no ABSOLUTE metric exceeded |r| = 0.25 against any axis), so differences in tumour content do not obviously account for the absence of an association, which remains unexplained.

Drug sensitivity in cell lines did not track the contrast. Of the 44 DepMap PDAC lines, 24 had drug-response data in GDSC2. Six MAPK-pathway inhibitors, chosen in advance, showed no association after correction, and all six correlations ran opposite to the predicted direction. Because the contrast includes BCL-2 family genes, we then tested nine BH3 mimetics, with nine cytotoxic agents as a secondary set, again with the drugs and expected direction fixed beforehand. Six of nine BH3 mimetics correlated in the predicted direction (median r = -0.034) but none survived correction; eight of nine cytotoxic agents did so (median r = -0.186), none nominally significant, and four of these are topoisomerase I inhibitors and therefore not independent (Supplementary Table 3). With 24 lines, moderate associations could have been missed, and viability assays count arrest as sensitivity, so they cannot separate the two outcomes the contrast is meant to distinguish.

## Discussion

Across 18 untreated patients, malignant cells were consistently weighted toward arrest-associated rather than apoptosis-associated expression, and the same direction appeared in an independent atlas scored identically. The shift was stable across scoring settings and was not reproduced by random gene sets matched for size and detection. Where the dividing line is drawn determines the percentage, so the finding is the direction of the shift and its consistency between patients rather than the figure of 86.3%.

That established tumour cells lean away from apoptosis is not in itself surprising; resistance to cell death is a defining feature of cancer, and cells in a resected tumour have already survived transformation. The useful part is that the extent of this bias can be measured in each cell, differs between patients and tracks tumour subtype. Three findings make that quantity worth measuring. In an inducible mouse model, a dormant subpopulation survived removal of oncogenic KRAS and drove relapse, depending on oxidative phosphorylation rather than glycolysis (Viale et al., 2014). In RAS-mutant cancers generally, MEK inhibition mostly halts growth rather than killing cells, and adding a BCL-XL inhibitor converts that stasis into regression, with BCL-XL identified by synthetic lethal screening as the main brake on apoptosis after MEK inhibition (Corcoran et al., 2013). Direct KRAS(G12D) inhibition is more potent and does induce apoptosis in mouse PDAC (Kemp et al., 2023), although durable regression still requires CD8 T cells (Mahadevan et al., 2023). Across these systems the recurring theme is that blocking RAS signalling does not by itself guarantee cell death, and that whether it does depends on how close the cell already sits to the apoptotic threshold.

The subtype association provides independent biological context for that interpretation. Tumours with higher basal-like scores contained fewer arrest-side cells, matching preclinical evidence that a classical epithelial state confers acute resistance to KRAS inhibition while basal-like and mesenchymal states respond better (Singhal et al., 2024; Dilly et al., 2024). The two literatures are independent: the subtype classifier was built from bulk expression to predict prognosis (Moffitt et al., 2015), and the modules here were curated from apoptosis and cell-cycle biology, yet they align in the direction that treatment response would predict. The association is also instructive where it fails. Basal-like tumours carry a worse prognosis but contain proportionally fewer arrest-associated cells, so the contrast is not a measure of aggressiveness, and it should not be expected to behave like one; this is consistent with the absence of a survival association in bulk data. Within tumours this relationship was not apparent: cells with higher scores for either Moffitt signature were similarly likely to fall on the arrest-associated side. The patient-level association may therefore reflect broader differences in tumour composition or biological context rather than a consistent within-tumour relationship between subtype and the contrast. With 18 patients and a wide confidence interval, this needs confirmation in a larger cohort.

TGM2 is the single gene the analysis points to, and its known biology in PDAC fits the side of the contrast on which it is enriched. TG2 is overexpressed in most PDAC tumours and cell lines, is associated with gemcitabine resistance and nodal metastasis, and constitutively activates focal adhesion kinase and PI3K-AKT survival signalling (Verma et al., 2006). Matrix-bound TG2 protects cells from detachment-induced apoptosis (Verderio et al., 2003), it participates in latent TGF-beta activation (Lockhart-Cairns et al., 2022; Eckert et al., 2014), and tumour-cell TG2 suppresses CD8 T cell responses in PDAC by promoting secretion of immunosuppressive cytokines (Lahusen et al., 2025). The enrichment of TGM2 among cells with relatively greater arrest-associated expression is consistent with reported roles for TG2 in promoting cell survival and resistance to apoptosis; the association also recurred across two platforms. Three limits apply. The effect is small, TGM2 came from an exploratory ligand-receptor screen rather than a pre-specified test, and the predicted TGM2-ITGA4 interaction has no protein-level or functional support here. TGM2 is therefore best read as a candidate marker of the arrest-associated state, and possibly a contributor to it, rather than an established mechanism.

Three further results define the boundaries of the finding. In tissue, suppressive and effector immune programmes followed the same gradient away from tumour epithelium, which fits the compartmental organisation of PDAC stroma, where regional immune composition follows fibroblast state rather than proximity to tumour cells (Grunwald et al., 2021). These spatial data do not establish a suppressive niche built by arrest-side tumour cells or by TGM2, and they illustrate the value of the effector control: without it, the Treg and checkpoint gradients alone could have been read as evidence of spatially organised suppression. In cell lines, the contrast did not track drug sensitivity, but the available viability assays count arrest and death alike and therefore cannot test a measure designed to separate them. A complementary functional approach is BH3 profiling, which tests how readily mitochondria undergo apoptosis. Both pre-treatment mitochondrial priming and its change during drug exposure can predict cytotoxic response where expression signatures do not (Ni Chonghaile et al., 2011; Montero et al., 2015). The contrast may capture transcriptional features related to apoptotic susceptibility, but whether it reflects functional mitochondrial priming remains unknown. This, together with the inability of conventional viability assays to separate arrest from cell death, may contribute to the absence of a drug-sensitivity association. In bulk tumours there was no clear association with survival. That is consistent with the possibility that the contrast measures a feature of cell state rather than prognosis, although the absence of an association remains unexplained.

Several limits bear on how far these data can be taken. The study measures transcript abundance, which constrains but does not determine whether a cell dies: apoptotic commitment also depends on protein levels, post-translational control and mitochondrial priming, and these can diverge from mRNA. Both cohorts are small surgical series of untreated, resectable tumours, whereas RAS inhibitors are used in advanced, previously treated disease, and no patient received a RAS inhibitor, so the trial that motivated the question is context rather than something these data explain. The modules are curated from established biology, which keeps them interpretable but means any PDAC-specific apoptotic programme would be missed. Single-nucleus and single-cell platforms also differ in transcript capture, which may contribute to the lower prevalence in the replication cohort, and dissociation itself can induce stress-response genes in scRNA-seq (van den Brink et al., 2017), although the direction of the shift was consistent despite these differences.

The central assumption can be tested directly. Patient-derived organoids profiled for the contrast, treated with RAS pathway inhibitors, and followed by live-cell imaging alongside BH3 profiling would establish whether cells on the arrest-associated side stop dividing rather than die, and whether the transcriptional contrast tracks the functional apoptotic threshold it is meant to approximate. If it does, the therapeutic question follows: whether lowering that threshold, as BCL-XL inhibition does after MEK inhibition, converts arrest into death preferentially in tumours weighted toward the arrest-associated side, with TGM2 the first candidate to test alongside it. Paired biopsies before and during RAS(ON) inhibitor treatment would then show whether the measurement carries clinical information. Until then, these data provide a reproducible description of where untreated PDAC cells sit between two programmes, a method for measuring it, and a clear statement of which of its implications remain untested.

## Ethical statement

Ethics approval and consent to participate: This study involved secondary analysis of publicly available, de-identified datasets. No new participants were recruited, no new human samples were collected, and no identifiable participant information was accessed. Ethics approval and participant-consent procedures for the original cohorts are described in the source publications (Hwang et al., 2022; Steele et al., 2020; Khaliq et al., 2024). The present secondary analysis did not involve contact with participants or access to identifiable data; no separate participant consent or study-specific ethics approval was obtained for this analysis. Consent for publication: Not applicable. The manuscript contains no identifiable individual-level information or participant images.

## Data availability statement

All source datasets analysed in this study are publicly available. DepMap 24Q2 data are available from the Broad Institute (https://depmap.org), and GDSC2 release 8.5 drug-response data are available from the Genomics of Drug Sensitivity in Cancer project (https://www.cancerrxgene.org/). GSE202051, GSE155698, GSE272362 and GSE269985 are available through the NCBI Gene Expression Omnibus; per-sample filtered count matrices for GSE155698 were obtained from the individual sample records. TCGA PAAD expression data are available from the NCI Genomic Data Commons (https://portal.gdc.cancer.gov), curated clinical annotations from the TCGA PAAD marker paper (Raphael et al., 2017), and ABSOLUTE tumour purity estimates from the GDC PanCanAtlas publication resources.

The R analysis scripts, project environment (including the renv lockfile), workflow documentation and supporting processed objects are publicly available in the GitHub repository (https://github.com/obuekweemeka/TGM2_PDAC). A versioned archive of the code is deposited in Zenodo (https://doi.org/10.5281/zenodo.23955811). The repository README describes the input-data retrieval steps, software requirements, analysis run order and the mapping between scripts and manuscript figures and tables. Original source datasets are not mirrored in the repository and should be obtained from the public repositories and accessions listed above. The scripts reproduce the reported statistics from the deposited processed objects.

## Acknowledgments

Not applicable.

## Author contributions (CRediT taxonomy)

Chukwuemeka A. Obuekwe: Conceptualization, Methodology, Formal Analysis, Visualization, Writing (Original Draft), Project Administration and Supervision. Blessing T. Ogunoye: Validation and Writing (Original Draft).

Aderinsola Adelaja: Validation, Methodology and Data Curation. Jude Ossai: Validation and Writing (Review & Editing).

Adegbesan A. Christopher: Writing, Methodology (Review & Editing) Adewunmi Akingbola: Formal Analysis and Data Curation.

