## Supplementary Figures and Table Legends for "Arrest-associated transcriptional programmes outweigh apoptosis-associated programmes in treatment-naive pancreatic ductal adenocarcinoma"

Supplementary Information

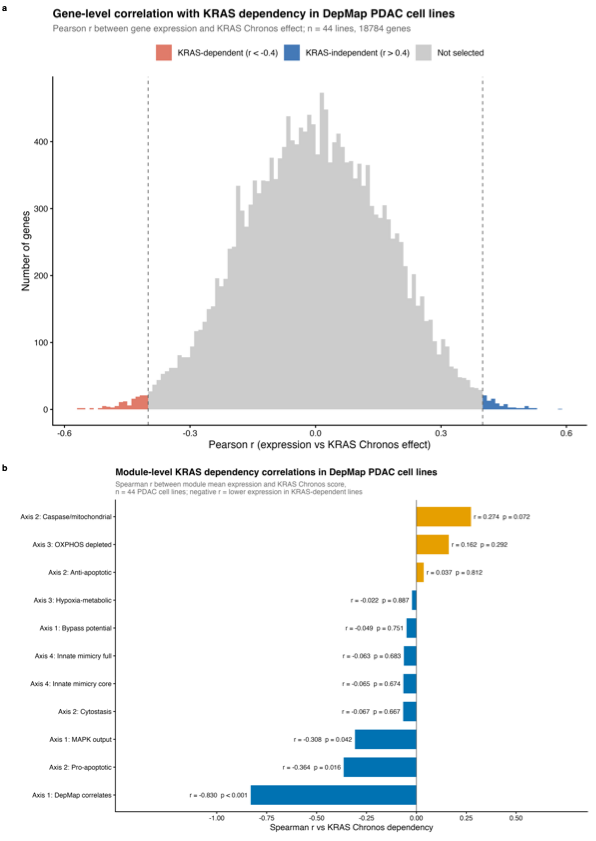

**Supplementary Figure 1.** Axis 1 and KRAS dependency in DepMap PDAC cell lines. a) Distribution of gene-level Pearson correlations between expression and KRAS Chronos gene effect across 18,784 genes in 44 PDAC lines; dashed lines mark the +/- 0.4 selection thresholds. The 30 most negative genes form the DepMap correlates module, which is circular by construction and not used as validating evidence. b) Module-level Spearman correlations against KRAS Chronos score for all 11 modules.

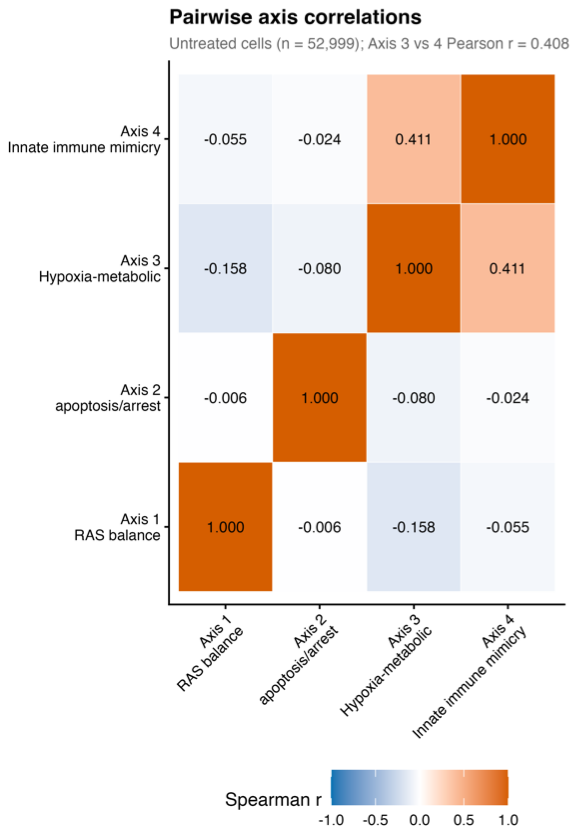

**Supplementary Figure 2.** Pairwise Spearman correlations between the four axis scores across untreated malignant cells. Axes 3 and 4 correlate at Pearson r = 0.408, below the 0.6 threshold specified in advance for merging them, so they were retained as separate axes.

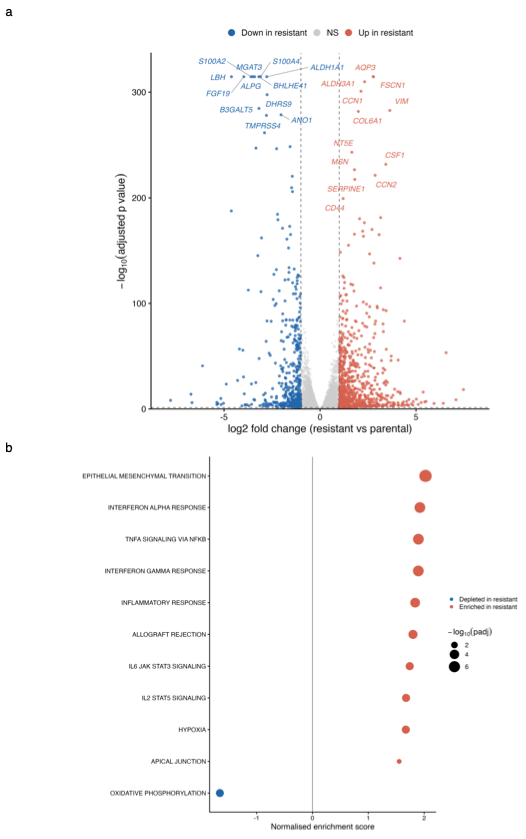

**Supplementary Figure 3.** Resistance-associated transcriptional changes and pathway enrichment in PDAC cell lines. a) Volcano plot of differential expression between resistant and parental cells in the Panc 02.03 contrast (GSE269985), with shrunken log2 fold-change estimates; dashed lines mark the significance and effect-size thresholds. This panel shows one cell line, not a combined two-line result. b) Hallmark gene sets significantly enriched or depleted in the same contrast (fgsea, FDR < 0.05), ordered by normalised enrichment score; point size indicates -log10 adjusted p-value.

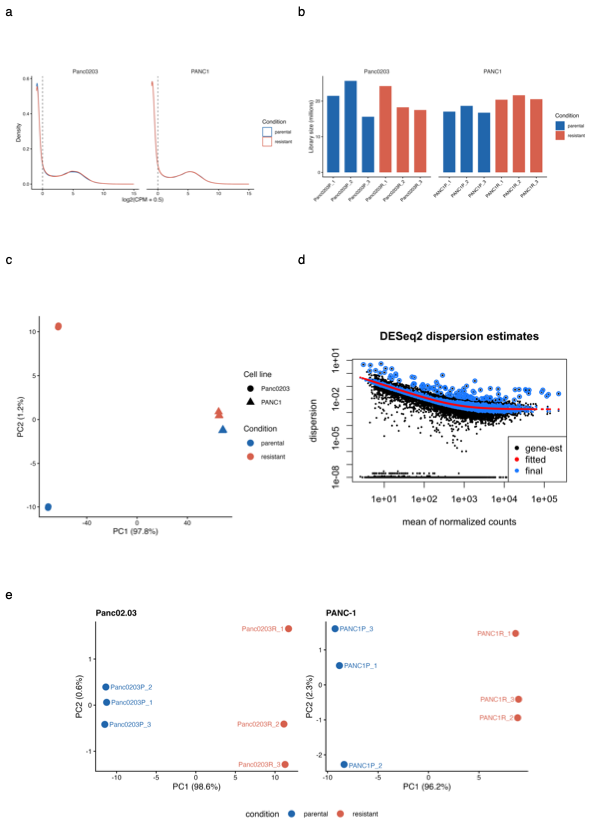

**Supplementary Figure 4.** Quality control for the resistance transcriptomics analysis (GSE269985). a) Distribution of log2(CPM + 0.5) per sample. b) Library sizes per sample. c) Principal component analysis of variance-stabilised counts across all 12 samples. d) DESeq2 dispersion estimates against mean normalised counts. e) Within-cell-line principal component analysis for Panc 02.03 and PANC-1 separately.

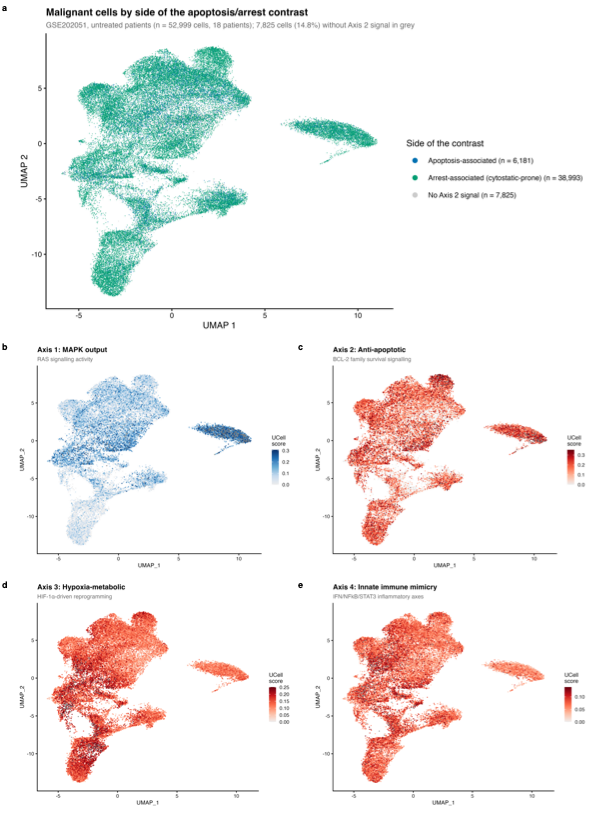

**Supplementary Figure 5.** Transcriptional landscape of treatment-naive malignant cells. a) UMAP of 52,999 malignant cells from 18 treatment-naive patients, coloured by side of the apoptosis/arrest contrast. Cells with a score of zero under the primary scoring procedure (7,825; 14.8%) are shown in grey and excluded from prevalence estimates. b to e) The same embedding coloured by the UCell score of one representative module for each axis: MAPK output (Axis 1), anti-apoptotic (Axis 2), hypoxia-metabolic (Axis 3) and innate immune mimicry (Axis 4). The embedding is used for visualisation only; no reported statistic depends on it.

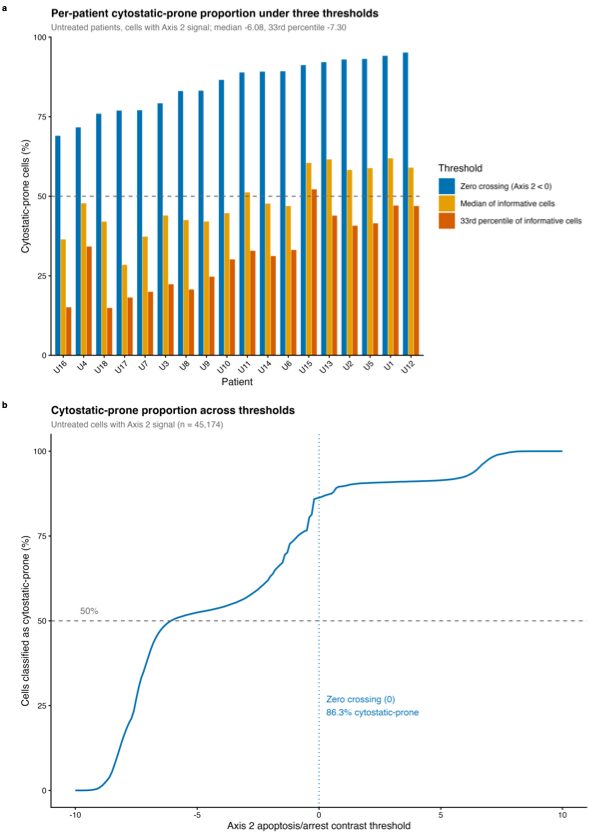

**Supplementary Figure 6.** Sensitivity of the classification to threshold choice. a) Per-patient proportions under three thresholds applied among cells with signal: the zero crossing, a median split, and a bottom-tertile split. Dashed line marks 50%. b) Proportion of informative cells on the arrest-associated side across the full range of thresholds. Pooling all cells the majority holds at any threshold above -6.0; a majority within every patient holds only above -1.0.

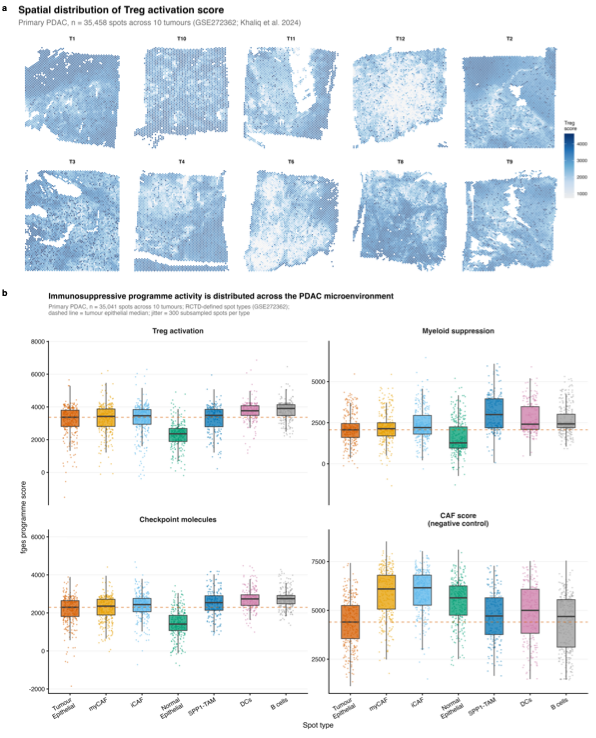

**Supplementary Figure 7.** Spatial data characteristics (GSE272362). a) Spatial distribution of the Treg activation score across 35,458 spots in 10 primary PDAC sections. b) Distribution of programme scores by dominant RCTD spot type across 35,041 spots, including a cancer-associated fibroblast score as a negative control. Scores are the precomputed fges values supplied with the published object.

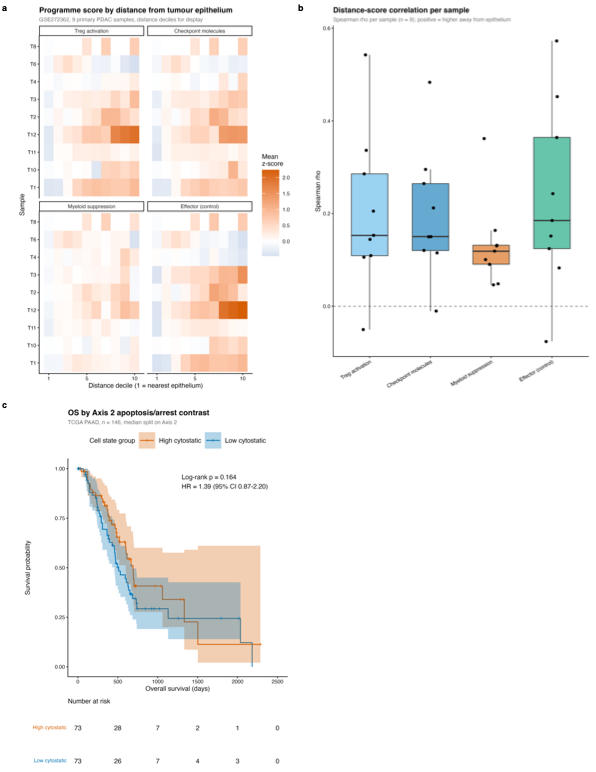

**Supplementary Figure 8.** Spatial organisation of immune programmes and bulk survival. a) Programme scores by distance decile from tumour epithelium across nine primary PDAC Visium sections, z-scored within section. b) Per-section Spearman correlation between distance from tumour epithelium and each programme score; all four programmes, including the effector control, increase with distance. c) Overall survival in TCGA PAAD by median split of the contrast score (n = 146; both Cox Wald and log-rank p-values reported in Results).

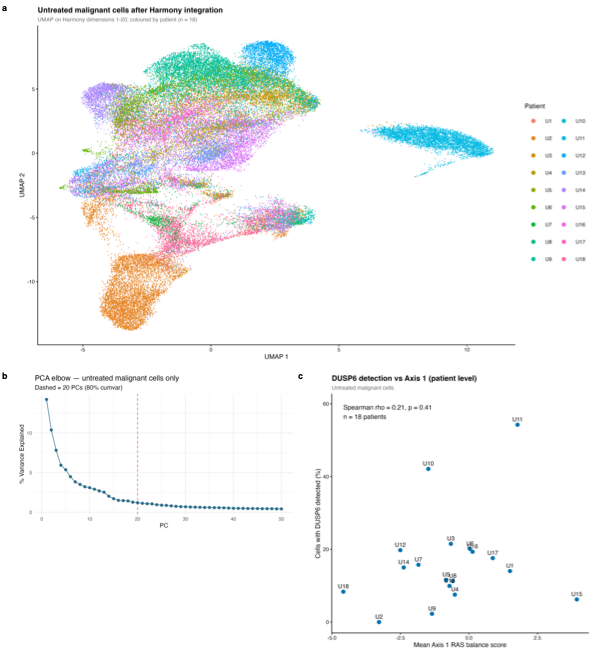

**Supplementary Figure 9.** Integration and axis quality control. a) UMAP of untreated malignant cells after Harmony integration, coloured by patient; cells remain partly separated by patient, as expected for clonal tumour transcriptomes. b) Scree plot of variance explained by principal component, with the dashed line marking the 20 components used. c) Patient-level relationship between the percentage of cells with detectable DUSP6 and mean Axis 1 score; DUSP6 detection does not track Axis 1, so Axis 1 has no patient-level external validation in this cohort.

**Supplementary Table 1.** Sensitivity of the classification to the UCell maxRank cutoff. For each cutoff: the number and percentage of treatment-naive cells returning an informative score, the percentage of informative cells on the arrest-associated side, the equivalent count and percentage against all 52,999 cells, the percentage unclassifiable, and the per-patient median, range and number of patients exceeding 50%.

**Supplementary Table 2.** Gene-set specificity of the contrast. Percentage of cells on the arrest-associated side under detection-matched random gene sets: 1,000 size-equalised permutations (10 genes against 10) and, for comparison, permutations preserving the original module sizes (13 against 10), alongside the observed value across 100 random equalisation draws.

**Supplementary Table 3.** Association between the cell-line apoptosis/arrest contrast and GDSC2 drug sensitivity across matched PDAC cell lines. For every compound with at least ten matched lines: the number of lines, Spearman correlation and p-value against ln(IC50) and against area under the curve, and Benjamini-Hochberg adjusted p-values within each primary drug set: the MAPK-pathway set defined before the first test and the BH3-mimetic set defined before the second. Cytotoxic agents, the secondary set of the second test, are reported without adjustment.
